# Elongation of stigmatic papillae induced by heat stress is associated with disturbance of pollen attachment in *Arabidopsis thaliana*

**DOI:** 10.1101/2019.12.21.885640

**Authors:** Kazuma Katano, Takao Oi, Nobuhiro Suzuki

## Abstract

Heat stress can seriously impact on yield production and quality of crops. Many studies uncovered the molecular mechanisms that regulate heat stress responses in plants. Nevertheless, effects of heat stress on the morphology of plants were still not extensively studied. In this study, we observed the detailed morphological changes of reproductive organs in *Arabidopsis thaliana* caused by heat stress. Larger area of stigma, and shorter length of anthers, filaments and petals were observed in plants subjected to heat stress compared to those under controlled conditions. Scanning electron microscopy (SEM) observation showed that length of stigmatic papillae without pollens seemed to be longer than that with pollens. In addition, classification of stigmas based on pollen attachment patterns together with artificial pollination assay revealed that pollen attachment onto stigma was clearly decreased by heat stress, and indicated that heat induced elongation of stigmatic papillae might be associated with disturbance of pollen attachment onto stigma. Furthermore, histochemical staining experiments revealed that crosstalk between Ca^2+^ and NO derived from pollens and O_2_^−^ derived from stigma might be associated with morphological alteration of stigma.

## INTRODUCTION

Negative effects of abiotic stresses on development of reproductive organs have been reported in recent studies. For instance, drought resulted in abnormal anther development, lower pollen viability, reduced filament elongation, ovule abortion and failure of flowering in Arabidopsis (Su et al., 2013). In contrast, enhanced elongation of stigmatic papillae was observed under high humidity (Takeda et al., 2018). These studies indicated that water status in surrounding environments alter the developmental patterns of reproductive organs, leading to decrease in seed production. Heat stress was also shown to negatively affect seed production (Katano et al., 2018b; Suzuki et al., 2013; Tunc-Ozdemir et al., 2013). We can therefore expect that decrease in seed production caused by heat stress might be at least partially attributed to morphological alterations in reproductive organs. Indeed, it has been well known that heat stress dramatically impacts on development of anthers and pollens (Katano et al., 2018a; Rieu and Twell, 2017). However, effects of heat stress on development of other reproductive organs are still largely unknown. In addition, only few studies focused on detailed morphological alterations of reproductive organs caused by heat stress, despite the remarkable progress of researches uncovering molecular mechanisms that regulate response in reproductive organs to heat stress.

In this study, to dissect the morphological alteration of reproductive organs induced by heat stress, we investigated the size of each reproductive organ in Arabidopsis plants grown under controlled and heat stress conditions. We also observed effects of heat stress on accumulation and distribution of Ca^2+^, O_2_^−^ and nitric oxide (NO), key signaling molecules to regulate development of reproductive organs in plants.

## RESULTS AND DISCUSSION

To analyze morphological alterations in reproductive organs caused by heat stress, we observed buds and flowers of plants grown under controlled conditions or subjected to heat stress by dissecting microscope and SEM (Fig. 1). Observation of flowers by microscope (by the methods mentioned in Fig.S1) revealed that diameter and area of stigma in plants subjected to heat stress significantly increased compared to that grown under controlled conditions (Fig. 1A, B). In addition, elongation of stigmatic papillae was observed in flowers subjected to heat stress (Fig. 1 C-H, Fig. S2). Interestingly, a lot of pollens were attached to stigmatic papilla which seemed to be shrinked under controlled conditions (Fig.1C, red rectangle), although only few pollens were attached to stigmatic papillae in plants grown under heat stress (Fig. 1D, E). These results indicate that enlargement of stigma due to the elongation of stigmatic papillae might be mainly triggered by the decrease of pollen attachment onto stigma by heat stress. Anthers, filaments and petals were shortened in flowers subjected to heat stress when compared to that under controlled conditions (Fig. S2A-I, L, M, O). However, structure of anthers and pollens in flowers subjected to heat stress was comparable to that under controlled conditions (Fig. S3). In addition, no significant effects caused by heat stress were observed in other parameters (Fig. S2A-I, J, K, N). In contrast to flowers, heat stress did not clearly alter the morphology of reproductive organs including stigma diameter in buds, except for anther length (Fig. S4, Fig. S5). Even under the SEM, only slight elongation of stigmatic papillae was observed in buds subjected to heat stress when compared to that grown under controlled conditions (Fig. S5 A-C).

**Fig. 1.**
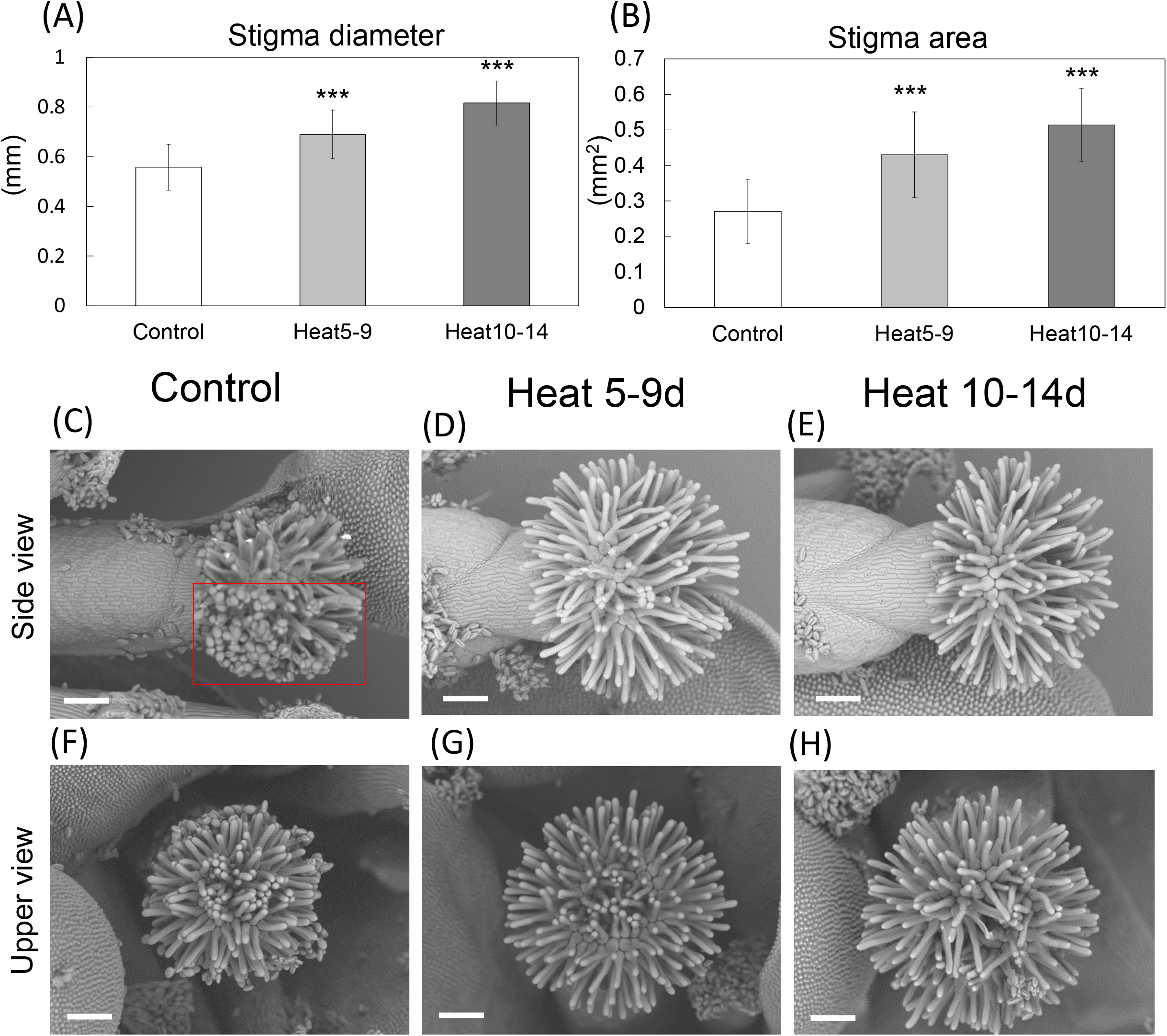
Enlargement of stigmas under heat stress. (A)-(B) Stigma diameter (A) and area (B) under controlled and heat stressed conditions. Bars in graphs indicate standard deviation. ***: Student *t*-test significant at p< 0.05, respectively compared to control (n=20-29). (C)-(E) Side view of stigmas grown under controlled conditions (C) or subjected to heat stress (D and E). (F)-(H) Top view of stigmas grown under controlled conditions (F) or subjected to heat stress (G and H). Photos were taken under the SEM. Red squares indicate the area of pollen attachment on stigma. White bars indicate 100mm.

Based on the SEM observation, we hypothesized that size of stigma might be determined by status of pollen attachment, and heat stress might disturb the pollen attachment onto stigma. To test this hypothesis, we classified the stigmas of flowers grown under controlled or heat stressed conditions into 4 different types based on pollen attachment patterns as indicated in Fig. S6 (Please also see “*Classification of pollen attachment patterns to stigma*” in “Materials and methods”). Fifty-eight percent of stigmas under controlled conditions showed type 1 pollen attachment pattern, and 14%, 7% and 21% of stigmas showed type 2, 3 and 4 pollen attachment pattern, respectively. On the other hand, only 7% of stigmas subjected to 5-9 days heat stress demonstrated type 1, and 4%, 25% and 64% of stigmas showed type 2, 3 and 4 pollen attachment pattern, respectively (Fig. 2A). Similar results were obtained in stigmas subjected to 10-14days heat stress; 10%, 16%, 16% and 58% of stigmas showed type1, 2, 3 and 4 pollen attachment patterns, respectively. These results clearly indicate that heat stress can disturb pollen attachment onto stigma. This disturbance of pollen attachment onto stigma might be at least partially due to the significantly longer distance between stigma and anthers under heat stress compared to that under controlled conditions (Fig. 2B). Shortening of filament length without alteration in style-ovary length might lengthen the distance between stigma and anthers. (Fig. S3K and M). We cannot ignore the possibility that damages on pollens might also affect development of stigma, because the shrinkage of anthers was observed in plants under heat stress conditions (Fig. S3L). We therefore analyzed pollen viability in buds of plants grown under controlled and heat stress conditions by Alexander staining (Fig. S7), and revealed that almost no pollens were alive in buds subjected to heat stress (Fig. S7A, B). Thus, low pollen viability could also affect development of stigmatic papillae under heat stress.

**Fig. 2.**
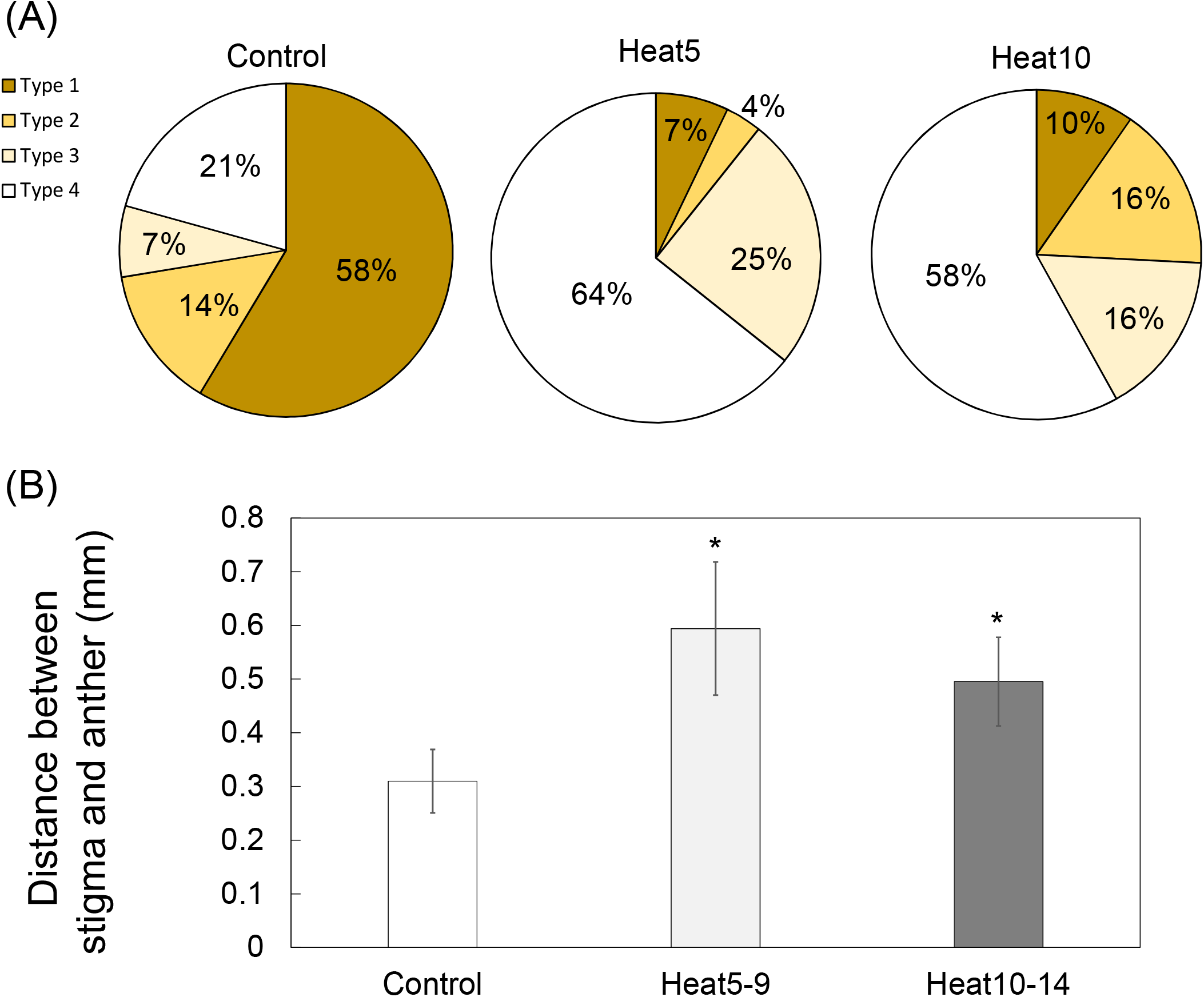
Analysis of pollen attachment patterns on stigma and distance between anthers and stigma. (A) Classification of stigmas into four types based on the pollen attachment patterns (refer to Fig S5). Proportion of each type of stigma relative to total number of stigmas tested in this study was calculated. (B) Distance between anthers and stigma grown under controlled or heat stressed condition. Bars indicates Standard error. *; Student *t*-test significant at p<0.05 compared to control (n=13-21).

To investigate the effect of pollen attachment on the size of stigma, we compared the diameter of stigma with or without artificial pollination (Fig. 3). The similar experiment was also conducted under heat stressed conditions to test if heat stress itself can cause enlargement of stigma without pollen attachment. Widened diameter of stigma without pollens was observed both under controlled and heat stressed conditions (Fig. 3A-C). On the other hand, shrinkage of stigma with pollens was observed under controlled and heat stressed conditions at 24h and 48h after artificial pollination (Fig. 3A-C). These results clearly demonstrate that enlargement of stigma was induced by disturbance of pollen attachment, not by heat stress. In addition, attachment of dead pollens did not induce shrinkage of stigmatic papillae (Fig.3 A-C). Furthermore, physical stimuli might not cause shrinkage of stigma, because attachment of silver nano particles did not clearly alter the size of stigma (Fig.3A-C). These results suggest that attachment of living pollens onto stigma might be required for the shrinkage of stigmatic papillae. Moreover, similar experiment was conducted using two other types of Arabidopsis (cv. Landsberg erecta and Wassilewskija), and similar results were obtained (Fig. S8A, B). These results indicate that the morphological alteration by pollen attachment onto stigma might be conserved process among the Arabidopsis varieties.

**Fig. 3.**
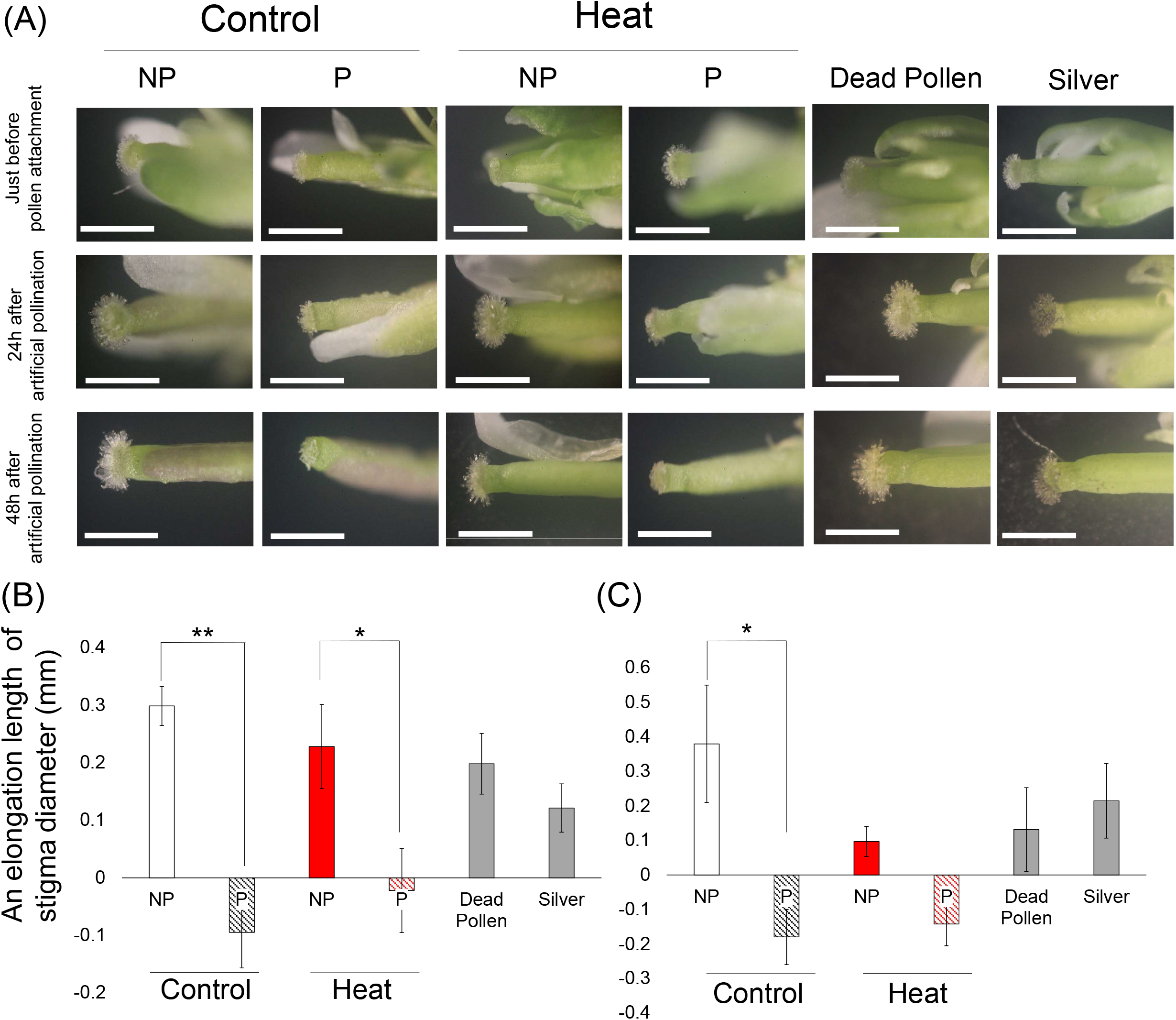
Effects of artificial pollination on stigma diameter under controlled and heat stressed conditions. (A) Photographs of stigmas before and after artificial pollination. NP; No pollens on stigma P; pollens on stigma. Silver particles and dead pollens also attached on stigma using as mechanical control. Bars indicate 1mm. (B) An elongation length of stigma diameter 24h after artificial pollination and (C) 48h after artificial pollination compared to that before pollination under controlled or heat stressed conditions. Bars indicates Standard error. * and **; Student *t*-test significant at p<0.05 and p<0.01, respectively compared to NP (n=5).

Previous studies revealed that crosstalk between Ca^2+^, O_2_^−^ and NO signaling regulates multiple processes of reproductive development (Katano et al., 2018a; Prado et al., 2004; Serrano et al., 2015; Traverso et al., 2013). We therefore analyzed the distribution and accumulation of Ca^2+^, O_2_^−^ and NO in flowers with poor or rich pollen attachments (i.e. type 4 or type 1 stigma, respectively; Fig S6). High accumulation of Ca^2+^ was detected in pollens, but, not in stigmatic papillae of flowers with type 4 stigma under controlled conditions (Fig. 4A-C). However, high accumulation of Ca^2+^ was observed in stigmatic papillae and pollens attached on it in flowers with type 1 stigma under controlled conditions (Fig. 4B, C). Similar results were obtained in flowers subjected to heat stress (Fig. 4G-I). Fluorescent intensity of Ca^2+^ level in type1 stigma is significantly higher than that in type 4 stigma both under controlled and heat stressed conditions (Fig. 4M). These results indicate that high accumulation of Ca^2+^ derived from pollens might be associated with shrinkage of stigmatic papillae. On the other hand, distribution of O_2_^−^ was only detected in stigma of flowers with type 4 stigma under controlled conditions. However, O_2_^−^ spread to style and ovary, but not accumulated in stigma in flowers with type 1 stigma under controlled conditions (Fig. S9A, B). This decrease of O_2_^−^ on stigma might be caused by process of pollen-pistil interaction (Zafra et al., 2016). Similar results were obtained in type 4 stigma subjected to heat stress, but O_2_^−^ spread was not observed in flowers with type 1 stigma subjected to heat stress (Fig. S9C, D). These results indicate that enlargement of stigmatic papillae might be associated with O_2_^−^ accumulation. In addition, NO detected in stigma with both type 4 and type 1 stigma under controlled and heat stressed conditions.

**Fig. 4.**
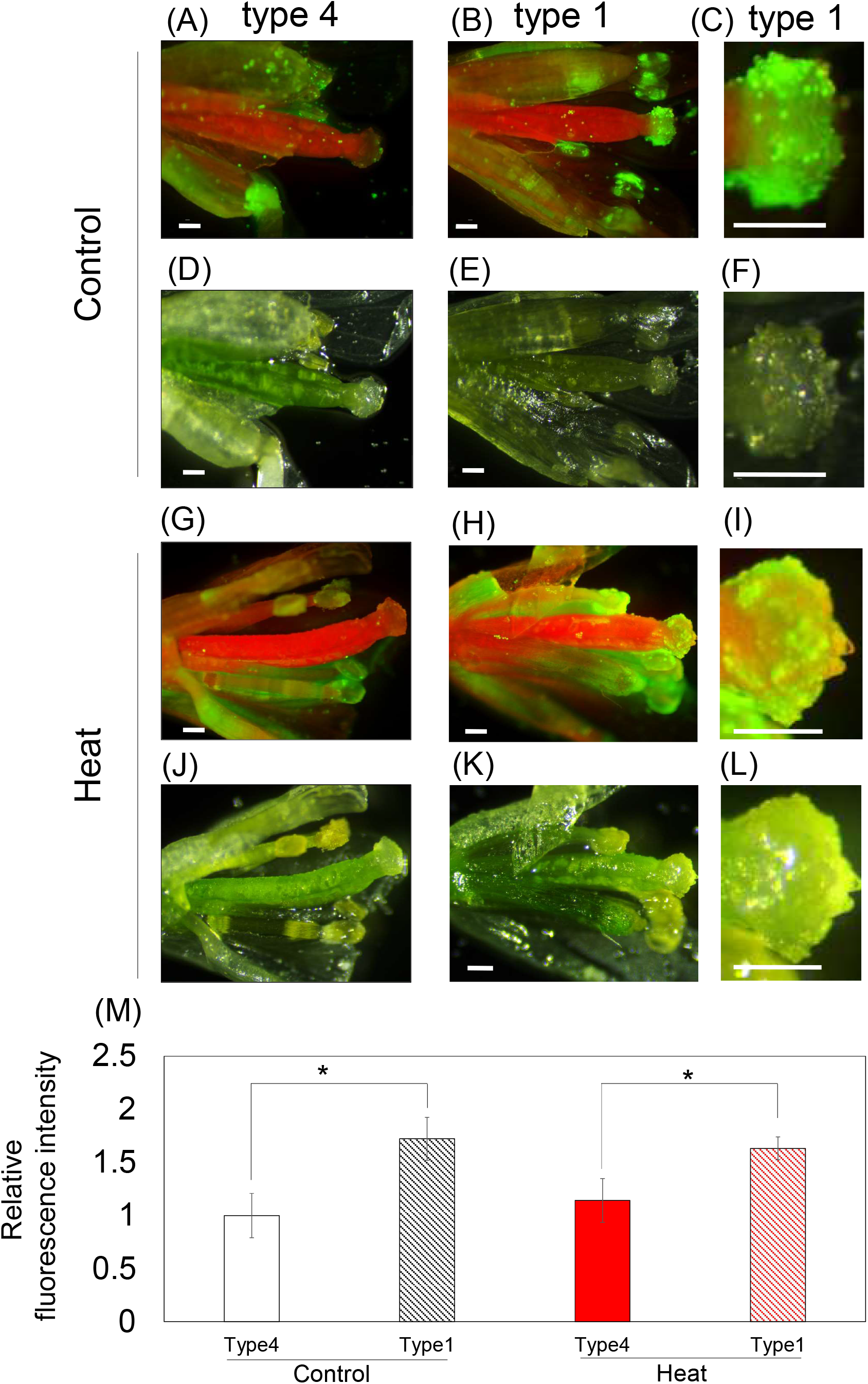
Distribution of Ca^2+^ in reproductive organs of *Arabidopsis* grown under controlled and heat stressed conditions. (A)-(L) Calcium green staining of reproductive organs in flowers with rich (type 1) or poor (type 4) pollen attachment under controlled (A)-(F) and heat stressed (G)-(L) conditions. (C), (F), (I) and (L) Enlarged figures of stigma in (B), (E), (H) and (K), respectively. (D)-(F) and (J)-(L) Bright field images. (M) Relative fluorescent intensity of Ca^2+^ level in stigma. The fluorescent intensity in stigmas relative to that in type 4 stigma under controlled conditions was calculated. Bars indicate standard error. *; Student *t*-test significant at p<0.05 compared to type 4 plants (n=5-7).

Furthermore, we also analyzed the distribution of Ca^2+^, O_2_^−^ and NO in buds under controlled and heat stressed conditions (Fig. S10). Ca^2+^ highly accumulated in anthers, but not in stigma under controlled and heat stressed conditions (Fig. S10A-D). In contrast to flowers, clear accumulation of O_2_^−^ and NO was not detected in reproductive organs both under controlled and heat stressed conditions (Fig. S10 E-J). These results also support the importance of crosstalk between these signals associated with pollen-pistil interaction for morphological alteration of stigma, because alteration in the size of stigma caused by heat stress was clearly observed in flowers in which pollen-pistil interaction proceeds, but not in buds (Fig. S4K, L; S10).

We discovered a novel phenomenon that pollen attachment onto stigma is clearly associated with stigma shrinkage, and disturbance of pollen attachment onto stigma by heat stress results in elongation of stigmatic papillae. A previous study demonstrated that high humidity condition induced morphological change in stigmatic papillae via regulation of ABA signaling (Takeda et al., 2018). In our study, enlargement of stigmatic papillae was observed under heat stress, suggesting that abiotic stress response mediated by ABA, an important stress response plant hormone could be involved in stigma enlargement. However, analysis employing artificial pollination in our study clearly showed significance of pollen attachment onto stigma, rather than heat stress, in stigma development (Fig. 4). This phenomenon that stigmatic enlargement when insufficient quant of pollen attached to stigma might be evolved in plants to increase the possibility of pollination success. Furthermore, numerous studies have strongly pointed out that attenuation of pollen production in anthers is associated with the sensitivity to heat stress (De Storme and Geelen, 2014; Rieu and Twell, 2017). Our results indicated that morphological alteration in stigma, as well as protection of pollen production, might be also an important factor to prevent the abortion of pollination during heat stress.

In this study, we also revealed that distribution of Ca^2+^ and ROS in reproductive organs altered depending on presence of pollens on stigma. We therefore hypothesized that crosstalk between Ca^2+^, ROS, and NO might be associated with morphological alteration of stigma depending on status of pollen attachment. Stigma without pollens showed high accumulation of O_2_^−^ in contrast to stigma with pollens under both conditions. Nevertheless, stigma without pollens showed almost no accumulation of Ca^2+^, although stigma with pollens showed high accumulation of Ca^2+^. It might be a unique phenomenon that can be observed on stigmas because Ca^2+^ signaling has been considered as inducer of RBOH dependent O_2_^−^ production. During pollination, however, it was an opposite event of the theory, which might be essential for proper pollen-pistil interaction (Kadota et al., 2015). This phenomenon might be involved in pollen recognition. In addition, during the pollen attachment on stigma, antioxidant substance in pollen coat was released into stigma via pollen hydration induced by NO-ROS redox signals (Katano et al., 2018a; Traverso et al., 2013). As for stigma with pollens, Ca^2+^ accumulated in pollens and clearly increased in stigma, suggesting that high accumulation of Ca^2+^ might be caused by activation of calcium permeable channel via NO derived from pollen (Besson-Bard et al., 2008; Clementi, 1998). Furthermore, O_2_^−^ and NO was not detected in each reproductive organs during bud stage under controlled and heat stress conditions. Considering only slight enlargement of stigma during bud stage, these results also support the importance of these signals for morphological alteration of stigmatic papillae. This crosstalk between Ca^2+^ derived from pollens and O_2_^−^ derived from stigma might be involved in stigma enlargement during poor pollination and shrinkage during rich pollination (Fig. S11). Indeed, involvement of Ca^2+^ that might function as a signaling molecule in tip growth of cells has been proposed in previous studies. For example, Ca^2+^ gradient with high accumulation in the tip was shown to be required for growth of pollen tubes and root hairs (Bibikova et al., 1997; Cardenas et al., 2008). However, Li et al., demonstrated that high concentration of extracellular Ca^2+^ inhibit the elongation of pollen tubes (Li et al., 1999). It is necessary to further elucidate how these signaling molecules including Ca^2+^ inhibit growth of stigmatic papillae cells in future studies.

Our results suggested that pollen attachment to stigma is a quite important factor for morphological alteration of stigmatic papillae. However, it is still unclear what are the factors in pollen associated with morphological alteration of stigma. One possibility is a pollen coat because pollen coat is a primal portion which directly attached to stigmatic papillae during pollination. Further studies will be required to elucidate significance of pollen coat in morphological alteration of stigmatic papillae.

## Materials and Methods

### Plant materials and growth conditions

*Arabidopsis thaliana* plants (cv. Columbia-0) were grown under controlled conditions (21°C, 16h light cycle, 50μmols^−1^m^−1^). Plants were grown on peat pellets (jiffy-7; http://www.jiffygroup.com/) using a growth chamber (LH-241SP, NK system, Tokyo, Japan). For the artificial pollination assay, two additional accessions of Arabidopsis (cv. Landsberg erecta and Wassilewskija) were also used.

### Stress treatment

For analysis of length of inflorescence stem, and number of buds, flowers and siliques, 25-day-old plants grown on peat pellets as described above were transferred to a growth chamber (LH-241SP, NK system, Tokyo, Japan) with the following temperature cycle; 06:00a.m.-09:00a.m., 21°C, 09:00a.m.-10:00a.m., 25°C, 10:00a.m.-12:00p.m., 40°C, 12:00p.m.-13:00p.m., 25°C, 13:00-06:00a.m., 21°C. Plants were then grown for additional 20 days under these temperature conditions. The 16h light period was imposed from 06:00a.m. to 10:00p.m. Control 25-day-old plants were maintained in parallel under controlled growth conditions for additional 20 days.

For analysis of morphological alterations in each reproductive organ, 30-day-old plants grown on peat pellets were subjected to heat stress with the same temperature cycle as described above or maintained under controlled conditions for 5-9 or 10-14 days. Buds and flowers were then collected from these plants and analyzed in size of each reproductive organ as described below.

### Observation of growth characteristics

Length of inflorescence stem, and number of buds, flowers and siliques of the plants grown under controlled conditions or subjected to heat stress were scored every two days during 20-day heat period.

Each reproductive organ was observed in buds and flowers collected from plants grown under controlled conditions, or subjected to heat stress for 5-9 or 10-14 days as described above using a stereoscopic light microscope (SZX12, Olympus、Japan) and a scanning electron microscope (SEM, TM3030, Hitachi, Japan). One to two petals and sepals were removed from buds and flowers in stage 11-12 and stage 13-15, respectively (Smyth et al., 1990). Buds and flowers were photographed before and after the removal of petals or sepals. Stigma area, stigma diameter, and length of flowers, style-ovary, anther, filament, petals and sepals as described in Fig. S1 were then measured on pictures using Image J. These parameters were measured in 8-20 flowers. Anther length and filament length were collected from 3-4 anthers and filaments per flower. Distance between stigma and anthers as described Fig. S1 were also measured on pictures using Image J. This parameter was analyzed by measuring distance between stigma and two anthers which were nearest to stigma.

Fine structure of stigma, anthers and pollens in buds and flowers corresponds to stage 11-12 and 13-15 (Smyth et al., 1990) were observed under SEM. Flowers collected from plants grown under the controlled conditions or subjected to heat stress were mounted on a stub with adhesive carbon tape, and transferred directly to specimen chamber of the SEM in low vacuum condition.

### Classification of pollen attachment patterns to stigma

Pattern of pollen attachment on stigma was analyzed in 28-31 flowers randomly collected from 10 plants grown under controlled conditions or subjected to heat stress. Flowers collected were photographed under light microscope and stigmas of these flowers were classified into four types based on the patterns of pollen attachment (Fig. S2). Type 1 indicates stigma almost completely covered with pollens. Type 2 indicates stigma attached with pollens on approximately half area. Type 3 indicates stigma attached by pollens only on an edge of stigma. Type 4 indicates stigma with no pollens. Then, proportion of each type of stigma under controlled or heat stressed conditions was calculated.

### Artificial pollination assay

All buds and flowers except for one biggest bud corresponding to stage 12 were removed from 30-day-old plants grown under controlled conditions as described above. Anthers were then removed from this remaining bud and pollens from other plants (Columbia-0) were attached onto the stigma of this remaining bud. One hour after artificial pollination, plants were subjected to moderate heat stress (35°C) for 1h. In this experiment, moderate heat stress was applied to buds, because heat stress (40°C, 2h) employed in the analyses of growth characteristics as described above was too strong for buds without sepals and petals. Stigmas were photographed at 0, 24h and 48h after exposure to heat stress and measured in diameter using Image J. Elongation length of stigma diameter was calculated by subtracting diameter before artificial pollination from that at 24h or 48h after artificial pollination. Stigma without artificial pollination were also photographed at the same time points in parallel and analyzed in elongation length of stigma diameter. For mechanical control, silver particles put on the stigma at the same timing of pollen attachment. Furthermore, Dead pollen also put on stigma as mechanical control. Dead pollen was using the anthers whose plants were grown under severe heat stress condition (42°C, 16h).

### Histochemical staining

*In situ* detection of O_2_^−^, Ca^2+^ and NO using nitro blue tetrazolium (NBT), Calcium green solution and Diaminofluorescein-2 (DAF-2), respectively were performed as previously described (Farnese et al., 2017; Iwano et al., 2014; Xing et al., 2013). Flowers with type1 and type 4 stigma were detached from plants grown under controlled conditions or subjected to heat stress. For O_2_^−^ detection, flowers were submerged in NBT solution (1 mg ml^−1^ NBT plus 10mM NaN3 solution in 10mM potassium phosphate buffer pH7.8) and incubated for 40 min at room temperature. Then, flowers were boiled in 95% ethanol for 15 min to completely remove the chlorophyll and stored in 60% glycerol. Flowers were observed by light microscopy and photographed. For Ca^2+^ detection, a drop of 10 μM Calcium Green solution containing 0.005% Tween 20 was put on flowers. After air dried for 1h, fluorescent signals in flowers were detected by fluorescence microscopy (LeicaMZ10 F) using 480/40nm excitation and 510 nm emission filter (GFP2 Plus). For NO detection, flowers were submerged in 10 μM of DAF-2 solution in the dark for 20 min at room temperature. Fluorescent signals were detected by fluorescence microscope (LeicaMZ10 F) using 480/40 nm excitation and 510 nm emission filter (GFP2 Plus).

### Alexander staining

Pollen staining with the Alexander dye was performed as previously described (Zhou et al., 2013). Anthers were collected from bud corresponds to stage 11-12 (Smyth et al., 1990) in plants grown under controlled conditions or subjected to heat stress. Anthers were then fixed with a 6:3:1 (v:v:v) of ethanol: chloroform: acetic acid for 2 hours. Following the fixation, anthers were put on a microscope slide and 2 or 3 drops of Alexander solution were applied. Slides were slowly over an alcohol burner in a fume hood until the stain solution is near boiling (~30 seconds). Heating can be adjusted by briefly moving the slide in and out of the flame. The cover-slip can be sealed and slides were examined using a microscope and taking photos of anthers.

## Acknowledgement

The authors would like to thank Dr. Makoto T. Fujiwara for experimental support. This work was supported by Sophia University.

